# Anthropogenic nest material use correlates with human landscape modifications in a global sample of birds

**DOI:** 10.1101/2023.06.16.545374

**Authors:** Catherine Sheard, Lucy Stott, Sally E. Street, Susan D. Healy, Shoko Sugasawa, Kevin N. Lala

**Affiliations:** School of Earth Sciences, University of Bristol, Life Sciences Building, 24 Tyndall Ave., Bristol BS8 1TQ, United Kingdom; Department of Anthropology, Durham University, Dawson Building, South Road, Durham, DH1 3LE, United Kingdom; School of Biology, University of St Andrews, Harold Mitchell Building, St Andrews KY16 9TJ, United Kingdom

**Keywords:** bird nests, nest material, artificial material, plastics, conservation

## Abstract

As humans increasingly modify the natural world, many animals have responded by changing their behaviour. Predicting the extent of these responses is a key step in conserving these species. For example, the tendency for some species of birds to incorporate anthropogenic items – particularly plastic material – into their nests is of increasing concern, as in some cases this behaviour has harmful effects on both adults and young. Studies of this phenomenon, however, have to date been limited in geographic and taxonomic scope. To investigate the global correlates of anthropogenic (including plastic) nest material use, we used Bayesian phylogenetic mixed models and a dataset of recorded nest materials in 6,147 species of birds. We find that after controlling for research effort, anthropogenic nest material use is correlated with proximity to human landscape modification, synanthropic (artificial) nesting locations, breeding environment, and the number of materials that has been recorded within the species’ nest. We also demonstrate that anthropogenic nest material use is unrelated to body mass, range size, or conservation status. These results indicate that anthropogenic materials are more likely to be included in nests when they are more readily available, as well as potentially by species who have more flexibility in nest material choice.

## Introduction

The Earth’s now over eight billion human inhabitants have left a significant mark upon the natural world (Crist et al., 2017; Venter et al., 2016). Many non-human animals (hereafter ‘animals’) currently live within or in close proximity to human-modified landscapes, and some have remarkable morphological or behavioural adaptations to such conditions (Alberti, 2015; McDonnell & Hahs, 2015; Palumbi, 2001). While a number of species thrive in anthropogenic settings (e.g., the success of urban red foxes, Plumer, 2014; or the persistence of threatened parrots within cities, Luna et al., 2018), others have experienced rapid population declines or local extinctions (McKinney, 2008). The ability to predict species’ response to urbanisation and other human modifications would thus improve our ability to protect and conserve species vulnerable to such changes (Dornelas et al., 2014; Marzluff, 2001; McDonald et al., 2008; McGill et al., 2015).

The nest is key to the reproductive success of nearly all bird species (Collias & Collias, 2014; Hansell, 2000). A well-constructed nest will protect the eggs and young from predators and environmental pressures (Deeming & Reynolds, 2016; Mainwaring & Hartley, 2013; Reid et al., 2000), and the materials used to construct a nest can reflect various physical and mechanical properties known or thought to contribute to offspring survival (Bailey et al., 2014; Bailey et al., 2016; Breen et al., 2021).

Some bird species, however, will incorporate anthropogenic (human-made) materials into their nests (reviewed in in Reynolds et al., 2019 and Jagiello et al., 2019). This phenomenon has ranged from a 1933 record of a pied crow (known then as *Corvus scapulatus*, now *Corvus albus*) placing various wire pieces into a 20-lb nest (Warren, 1933) to reports of plastic debris in 12% of nests from 14 northwest European seabird species (O’Hanlon et al., 2021). Sometimes anthropogenic materials appear to provide potential benefits, such as the reduction of ectoparasites in the nest due to the inclusion of cigarette butts by house sparrows (*Passer domesticus*) and house finches (*Carpodacus mexicanus*) (Suárez-Rodríguez et al., 2013). In other cases, this behaviour is known or presumed to be harmful; the incorporation of plastic into seabird nests, for example, puts individuals at higher risk of entanglement or ingestion (Gall & Thompson, 2015; Huin & Croxall, 1996; Montevecchi, 1991; O’Hanlon et al., 2021), and the same plastic string that strengthens great grey shrike (*Lanius excubitor*) nests in Poland also kills nestlings and breeding females (Antczak et al., 2010).

In most instances, however, the ecological and evolutionary causes and consequences of inter-specific variation in anthropogenic material nest usage have not yet been identified. In particular, most studies of anthropogenic nest material use have focused on a small number of species and/or have had limited geographic scope. For example, in addition to the aforementioned studies, anthropogenic nest materials have been studied in great tits (*Parus major*) and blue tits (*Cyanistes caeruleus*) in Warsaw, Poland (Jagiello et al., 2022); in satin bowerbirds (*Ptilonorhynchus violaceus*) at a single fieldsite in Australia (Borgia, 1985); in black kites (*Milvus migrans*) in Doñana National Park in Spain (Sergio et al., 2011); and in Chinese bulbuls (*Pycnonotus sinensis*) in Hangzhou, China (Wang et al., 2009). The extent to which the patterns of use found in these studies apply to all taxa and/or across all ecological settings remains unknown, thus hindering our ability to reach general conclusions as to the effects of the inclusion of anthropogenic material into bird nests.

Here, therefore, we present a global database of recorded nest materials (n = 6,147 species across 223 families), scored for the documented use of anthropogenic material. Due to the importance of plastic as a non-biodegrading and especially harmful material to birds (e.g., Townsend & Barker, 2014; O’Hanlon et al., 2021; Avery-Gomm et al., 2019), we also separately consider the use of plastic as a nest material. We seek to understand predictors of anthropogenic and plastic material use in the nests of bird species to extend this research beyond a handful of well-studied systems, and to explore potential conservation implications. To our knowledge, this is the first broad-scale assessment of anthropogenic nest material usage.

We tested four hypotheses for the incorporation of anthropogenic/plastic nest material in birds. First, we evaluated whether the anthropogenic/plastic material use is linked to greater flexibility in material choice (Hansell, 2007), indicated by a **greater number of different nest materials**, while controlling for research effort (Stutchbury & Morton, 2001; Xiao et al., 2017). Second, we assessed whether, at the global scale, species that living in closer proximity to **human-modified environments** will be more likely to include anthropogenic/plastic nest materials, as seen in some previous, population-level studies (e.g., Bond et al., 2012; Jagiello et al., 2019; O’Hanlon et al., 2021; Suárez-Rodríguez et al., 2013). Third, we tested whether species breeding in **different environments** will use anthropogenic/plastic nest materials in different proportions, as different biomes might have varying levels of nest material availability (Briggs & Deeming, 2016; Mennerat et al., 2009). We also investigated whether species that build **different types of nests** are more or less likely to incorporate anthropogenic/plastic nest materials, as different nest construction strategies have different energetic demands and thus may be more or less able to incorporate non-preferred material types (Mainwaring & Hartley, 2013). We also tested whether greater anthropogenic/plastic nest material use is correlated with species extinction risk, and we included body mass in all models, due to its myriad associations with other ecological and life history traits.

## Materials and methods

### Data collection

All available descriptions of nest materials were collated from three sources: the Handbook of the Birds of the World Alive (HBW; 2017-2018), Neotropical Birds Online (NBO; 2019-2020), and the Birds of North America Online (BNA; 2019-2021); note that subsequently all of these sources have been combined into a single resource, the Birds of the World (Billerman et al., 2022). These lists of materials were then scored as a binary trait for the presence of anthropogenic material, which included string, rope, fishing line, wire, aluminium foil, cloth, paper, rubbish/trash, concrete fragments, cellophane, etc. We included all materials manipulated by the bird and attached to the egg cup within these lists, including materials used as nest lining (though not including materials placed during the construction process but then removed prior to egg laying); this was primarily because the authors of these sources rarely differentiated among the various structural functions of nest materials, but also because we had no a priori hypotheses to test regarding this distinction.

We additionally separately scored these lists for the presence of plastic material specifically; note that this variable may be under-documented even within the context of these lists, as unspecified materials such as ‘rubbish’ maybe have contained plastic but could not with certainty be counted as plastic presence. For each species, we also recorded the total number of different nesting materials use (i.e., the number of distinct material types listed, as separated by a comma, the word ‘or’, or the word ‘and’); if a species had multiple entries across the three sources, the maximum number of materials per source was taken as the species value.

Nest structure and location were also scored based on HBW, NBO, and BNA entries and photographs. Structure was marked as presence-absence for no nest or a scrape (i.e., no constructed nest, but material used as liner); a platform; a cup; a dome (including multi-chambered dome-and-tube nests); and an excavated nest (including nests where a cavity is excavated and then a nest is constructed inside). Location was marked as presence-absence for an artificial location (e.g., nest boxes, telephone poles, house eaves, etc.); on the ground or touching water; inside an earthen or tree cavity; on or within rocks raised above the ground; and attached to vegetation (e.g., reeds, bushes, trees). Uncertainty in nest categorisation – either noted in the entry itself (e.g., ‘dubious record’) or due to coder interpretation (e.g., an unclear description or photograph) – was regarded as trait absence, and disagreement between sources was resolved in favour of trait presence. Subsets of the six coders met regularly to discuss questions and spot-check each other’s work, and approximately one third of entries were later checked by at least one of the two most experienced coders. All coders were followed a detailed data collection manual and had a formal biological background (one undergraduate student, one post-baccalaureate researcher, one Masters student, and three postdoctoral researchers).

Body mass and range size data were obtained from Sheard et al. (2020). Brain size data was taken from the compilation published in Hooper et al. (2022) and averaged for a single per-species value. IUCN 2020 Red List status was obtained where possible from BirdLife International (IUCN, 2022) on 2022-11-07 and scored from the IUCN Red List website (https://www.iucnredlist.org/) for the handful of species whose taxonomy could not be reconciled with BirdLife’s. To improve the accuracy of our parameter estimates, and as we had no *a priori* biological reason to distinguish these particular categories, we reclassified ‘endangered’ species include both endangered and critically endangered species; ‘extinct’ species includes species both extinct and extinct in the wild; and ‘threatened’ species include near threatened and vulnerable species. We excluded data-deficient species from analyses containing this conservation status variable.

Biome membership was scored by intersecting the 2018 BirdLife International range maps (BirdLife International, 2018) with the World Wildlife Fund (WWF) global terrestrial biome data (Olson et al., 2001) and taking as the biome identity for each species the biome with the greatest proportional intersection; birds with largely non-terrestrial ranges were scored as ‘seabirds’, and a few birds were hand-scored according to IUCN Red List habitat information and/or Birds of the World habitat information due to taxonomic mismatches or errors with the range maps. WWF biomes 1, 2, and 3 were considered ‘tropical forests’; biomes 4 and 5 were considered ‘temperate forests’; biomes 6 and 11 were combined into a single tundra-taiga category; biomes 7, 8, and 10 were considered ‘grasslands’; biome 12 was considered ‘Mediterranean’; biome 13 was considered ‘desert’; biome 14 was considered ‘mangroves’. As a measure of proximity to human landscape modification, a per-species score for human-footprint index (HFI, a global score from 0 to 100 on each terrestrial square kilometre measuring the human impact on the landscape, such as urbanisation, farmland, roads, etc.) was calculated by intersecting the Human Footprint Index (Wildlife Conservation Society, 2005) with a 1°x1° grid and finding the average value for each grid cell, multiplying these values as a vector across a presence-absence grid for the world’s birds based on the 2018 BirdLife International range maps, and then calculating the arithmetic mean for each species. Finally, as many of these variables are likely to co-vary with the amount of scientific investigation of each species’ ecology, we recorded the number of articles returned by a Web of Science search of each scientific name (2023-01-12 to 2023-01-23) as a proxy for research effort.

### Phylogenetic comparative methods

To test for the associations between anthropogenic/plastic nest material use and the proposed explanatory variables, we used Bayesian phylogenetic logistic regressions in the package *MCMCglmm* (Hadfield, 2010) in R version 4.1.3 (R Core Team, 2022). Anthropogenic and plastic nest material use were each considered as binary response variables; fixed effects were all included within a single model to reduce Type I error and to control for covariation between these possible predictors. Body mass was log-transformed prior to analysis, research effort was square-root transformed, and the continuous variables of material number, body mass, range size, research effort, and human-footprint index were each rescaled to have a mean of 0 and a variance of 1 to improve coefficient interpretability. A sample of 100 phylogenetic trees were obtained from the Hackett backbone of the Global Bird Tree (Jetz et al., 2012), trimmed to match the data from each model, and included as random effects within each model. Note that species missing any data (from the predictor variables or the phylogeny) were omitted from the analysis. Priors for the fixed effects were determined using Gelman priors (Gelman et al., 2008), with the prior for the phylogenetic variance set to *V* = 1^-10^ and *ν* = -1 and with the residual variance fixed to 1. For each of the two models (anthropogenic and plastic material use), an initial ‘dummy’ run was used to determine a start point on an arbitrary tree topology for 11,000 iterations, with a burn-in of 1,000 and a sampling rate of 10. We then looped across each of the 100 tree topologies for 30,000 iterations for each tree, with a burn-in of 10,000 and a sampling rate of 2,000, for a total of 10 stored iterations per tree. Model outputs were visually inspected to ensure convergence and proper mixing, and all effective sample sizes were greater than 200.

These models containing all potential correlates of anthropogenic and plastic material use contained fewer species (n = 4,237) than our total sample of all species with both nest material and phylogenetic information (n = 5,960). We therefore employed model selection procedures on both sets of response variables as sensitivity analyses, to verify that our results were robust to larger sample sizes. In brief, we sequentially compared the DIC fit between our main model and versions run with individual statistically non-significant fixed effects removed; at each iterative step, if any newly-accepted model could be performed on a larger sample of species, this model was re-run and this larger sample considered at the subsequent model selection step. We halted this procedure when either we found the best-fitting model according to DIC fit or when all statistically non-significant variables had been removed from the model.

Finally, to assess the relationship between relative brain size (a common proxy for noephilia and cognitive performance – see, e.g., Lefebvre et al., 2004; Sol et al., 2008; Sol et al., 2014; Sol et al., 2002) and plastic and anthropogenic material use, we ran a separate set of models on the reduced dataset for which we were able to obtain brain size data (n = 760 across all variables).

## Results

Of the 7,148 nest material entries that we were able to obtain (including several instances of multiple entries per species), 327 entries mentioned anthropogenic nest material (and 102 entries mentioned plastic material). According to the Birds of the World taxonomy, these entries combine to a count of 6,147 species, of which 291 (4.7%) were documented as building nests with anthropogenic material and 92 (1.5%) with plastic material (Figure 1). The orders with the highest proportion of anthropogenic material include the Coraciiformes (kingfishers, bee-eaters, motmots; 1 of 6 species, 17%), the Ciconiiformes (storks; 3 of 20 species, 15%), the Falconiformes (falcons and caracaras; 2 of 16 species, 13%), and the Suliformes (gannets, cormorants, frigatebirds; 6 of 48 species, 13%); the orders with the highest proportion of plastic material included the Suliformes (5 of 48 species; 10%), the Strigiformes (owls; 1 of 16 species, 6%), and the Pelecaniformes (pelicans, herons, ibises; 4 of 100 species, 4%).

**Figure 1:**
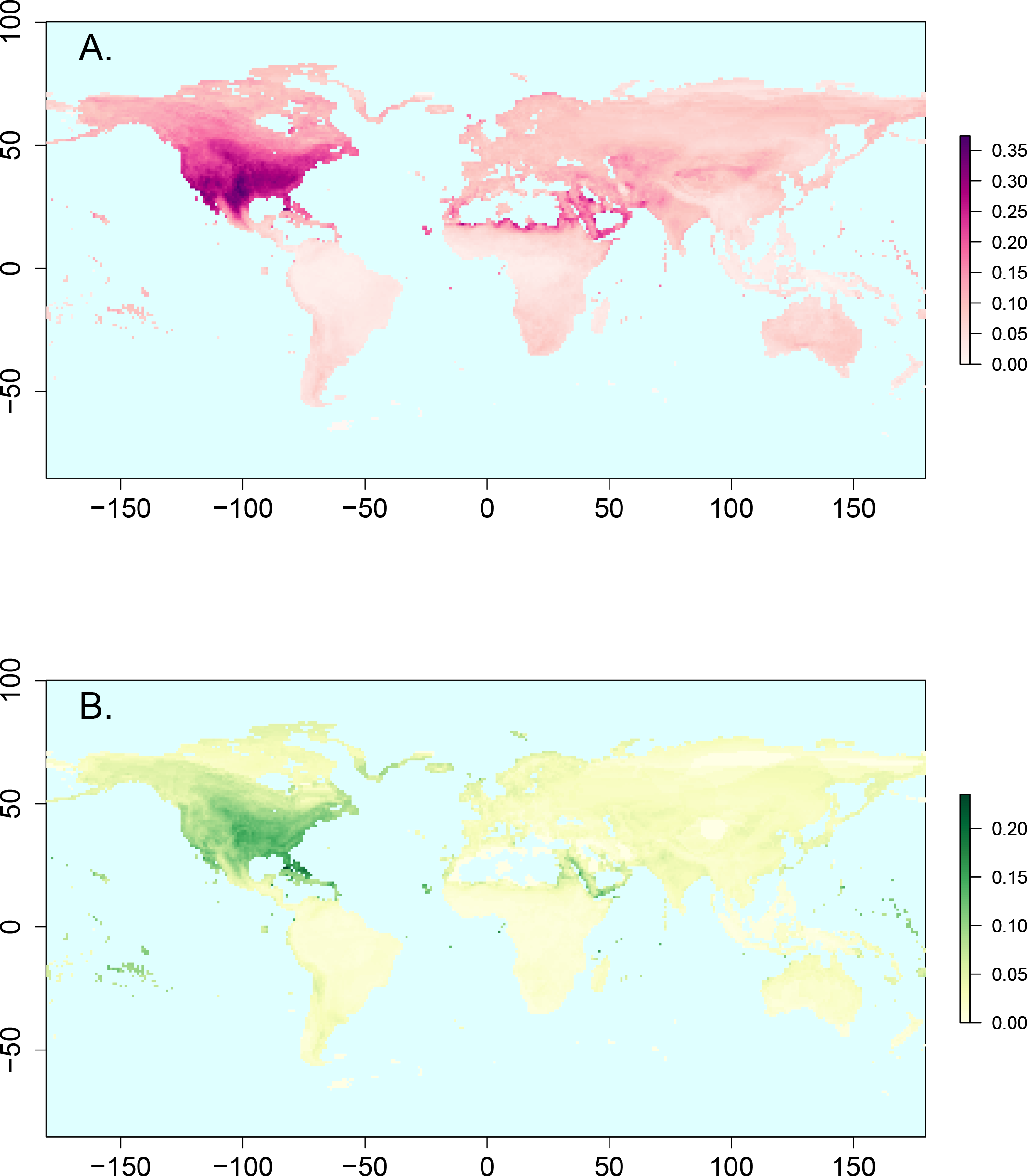
Global distribution of recorded anthropogenic and plastic nest material use. Darker colours indicate greater proportions of recorded anthropogenic (purple, panel A) and plastic (green, panel B) nest material use among the species breeding there, at the 1°x1° scale. Locations with fewer than 20 species with recorded nest material use have been omitted (e.g. interior Greenland, Sahara Desert).

Note that the BirdTree (Jetz et al., 2012) taxonomy, on which the phylogenetic models were based, contains fewer species than the Birds of the World, and thus the comparative models were based on at most 5,960 species, of which 282 were recorded to use anthropogenic material and 90 to use plastic material.

Globally, after controlling for a positive effect of research effort (*β* = 0.344, *pMCMC* = 0.004, Table 1, Figure 2), species were more likely to incorporate anthropogenic materials into their nests if they breed in more heavily human-modified landscapes (*β* = 0.479, *pMCMC* < 0.001), nest in synanthropic (human-modified) locations (*β* = 1.878, *pMCMC* < 0.001), and/or are documented as incorporating multiple more types of materials into their nests (*β* = 1.739, *pMCMC* < 0.001). The tendency to incorporate anthropogenic nest material also varies by biome membership; the inclusion of anthropogenic nest material is highest in deserts, lowest in tropical forests, and intermediate in grasslands, taiga/tundra, and temperate forests. There are no significant correlations between anthropogenic material use and body mass, nest structure, range size, or IUCN conservation status. The model selection procedure preferred this full model, despite the inclusion of non-significant variables (Table S1).

**Table 1:** Results of a Bayesian phylogenetic mixed model predicting interspecific variation in anthropogenic nest material use. Coefficients above 0 indicate a positive correlation with anthropogenic nest material use within a multivariate framework; coefficients below 0 indicate negative correlation. Conservation parameters are compared with a baseline of endangered species; breeding biome parameters are compared with a baseline of desert location. As many species nest in multiple structures and locations, these are included in the model as separate binary variables rather than as single categorical variables. Statistically significant coefficients are highlighted in grey. CI = credible interval. LC = least concern. Med. = Mediterranean. HFI = human-footprint index.

|  | coefficient | lower 95%<br>CI | upper 95%<br>CI | pMCMC |
| --- | --- | --- | --- | --- |
| Research Effort | 0.344 | 0.098 | 0.570 | 0.004 |
| # Materials | 1.739 | 1.478 | 2.081 | < 0.001 |
| Body Mass | -0.682 | -1.702 | 0.171 | 0.142 |
| Range Size | 0.035 | -0.205 | 0.266 | 0.760 |
| HFI | 0.479 | 0.196 | 0.774 | < 0.001 |
| Conservation - Extinct | -1.739 | -8.283 | 4.229 | 0.664 |
| Conservation - LC | -0.701 | -1.831 | 0.352 | 0.216 |
| Conservation - Threatened | -0.917 | -2.241 | 0.471 | 0.196 |
| Biome - Grasslands | -1.080 | -1.799 | -0.381 | 0.002 |
| Biome - Mangroves | -2.535 | -8.695 | 3.625 | 0.434 |
| Biome - Med. | -0.984 | -2.216 | 0.279 | 0.124 |
| Biome - Marine | -0.872 | -2.761 | 0.847 | 0.356 |
| Biome - Taiga/Tundra | -1.297 | -2.335 | -0.047 | 0.020 |
| Biome - Temperates | -1.591 | -2.487 | -0.773 | < 0.001 |
| Biome - Tropics | -2.318 | -3.083 | -1.531 | < 0.001 |
| Nest structure - scrape/none | -0.418 | -1.779 | 0.878 | 0.528 |
| Nest structure - cup | -0.767 | -2.007 | 0.409 | 0.192 |
| Nest structure - platform | -0.184 | -1.545 | 1.391 | 0.790 |
| Nest structure - dome | -0.644 | -2.005 | 0.553 | 0.316 |
| Nest structure - excavation | 0.033 | -1.514 | 1.701 | 0.970 |
| Nest location - artificial | 1.878 | 1.241 | 2.472 | < 0.001 |
| Nest location - ground/water | -0.124 | -0.818 | 0.618 | 0.734 |
| Nest location - cavity | -0.331 | -1.241 | 0.478 | 0.494 |
| Nest location - elevated rock | -0.207 | -0.983 | 0.547 | 0.580 |
| Nest location - vegetation | 0.070 | -0.823 | 0.912 | 0.870 |

**Figure 2:**
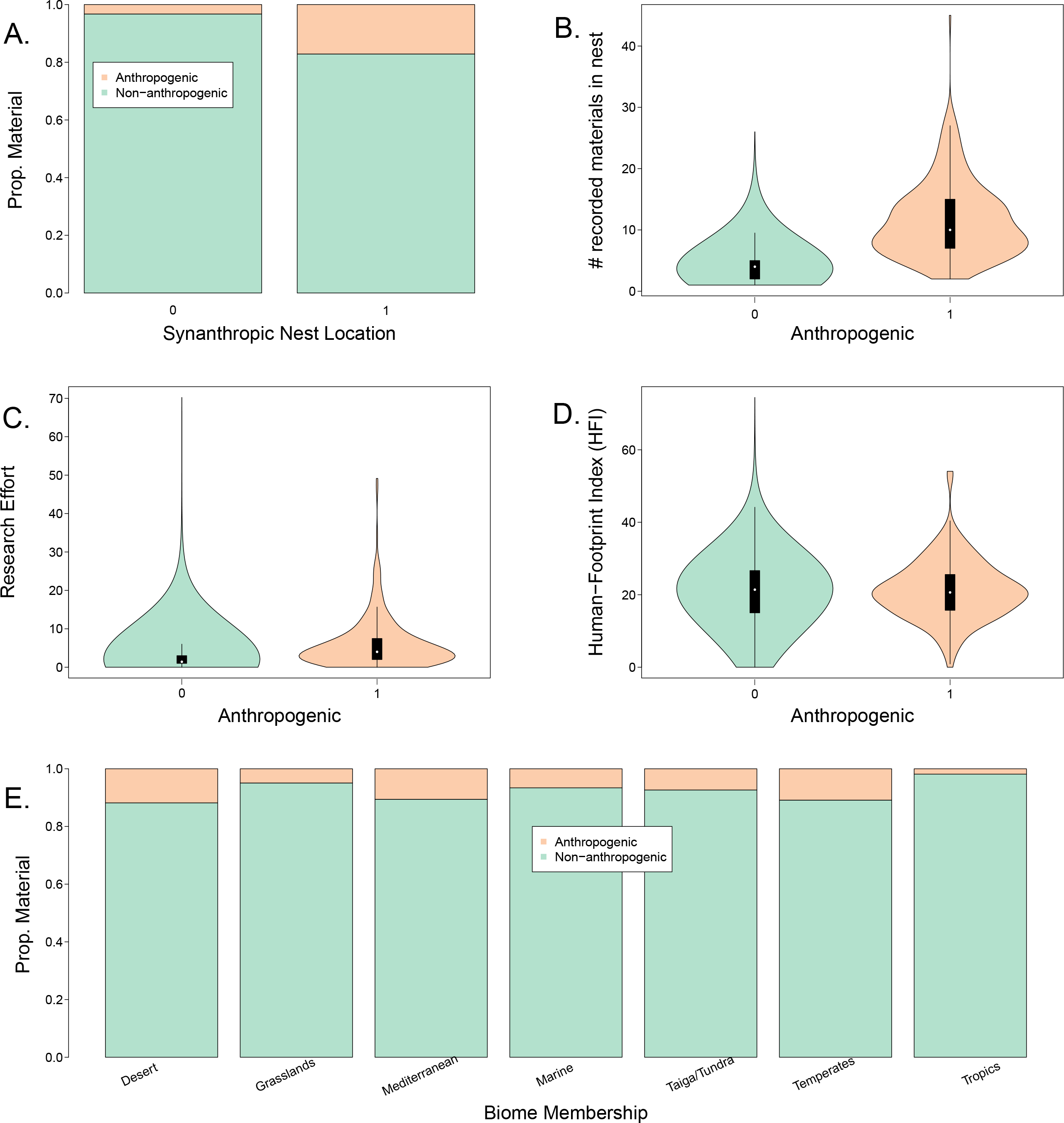
Ecological and synanthropic correlates of anthropogenic nest material use. Frequencies, uncorrected for phylogenetic signal or covariance with any other predictor, are presented at the species level between species that are (peach) and are not (green) known to use anthropogenic nest material and A. species that are and are not known to nest in synanthropic locations, B. the number of nest materials recorded for each species, C. ‘research effort’, here estimated by the number of papers indexed by the Web of Science about that species, D. the average human-footprint index for that species, and E. assigned biome membership. For ease of display in panel E, the eleven species that breed predominantly in mangroves have been omitted. All patterns displayed are statistically significant in the phylogenetically-corrected model after controlling for multiple co-variates (see Table 1). Prop. = proportion.

Bird species that use plastics in their nests also tended to breed in more heavily human-modified landscapes (*β* = 0.569, *pMCMC* = 0.004, Table 2, Figure 3), nest in synanthropic (human-modified) locations (*β* = 1.143, *pMCMC* = 0.004), and/or were recorded as incorporating more types of materials into their nests (*β* = 1.141, *pMCMC* < 0.001), after controlling for the positive effect of research effort (*β* = 0.482, *pMCMC* < 0.001). Species were also more likely to have been recorded as using plastic in their nest if they use materials as liner but do not construct full nest structures (i.e., were scored as using ‘scrapes’ or ‘no nest’; *β* = 1.644, *pMCMC* = 0.022), and plastic incorporation was less common in grasslands and in tropical forests than in other biomes. As with anthropogenic materials, plastic nest material use was apparently not related to body mass, range size, or conservation status. The model selection procedure again preferred this full model, despite the inclusion of non-significant variables (Table S2).

**Table 2:** Results of a Bayesian phylogenetic mixed model predicting interspecific variation in plastic nest material use. Coefficients above 0 indicate a positive correlation with plastic nest material use within a multivariate framework; coefficients below 0 indicate negative correlation. Conservation parameters are compared with a baseline of endangered species; breeding biome parameters are compared with a baseline of desert location. As many species nest in multiple structures and locations, these are included in the model as separate binary variables rather than as single categorical variables. Statistically significant coefficients are highlighted in grey. CI = credible interval. LC = least concern. Med. = Mediterranean. HFI = human-footprint index.

|  | coefficient | lower<br>95% CI | upper<br>95% CI | pMCMC |
| --- | --- | --- | --- | --- |
| Research Effort | 0.482 | 0.223 | 0.736 | < 0.001 |
| # Materials | 1.141 | 0.855 | 1.378 | < 0.001 |
| Body Mass | -0.960 | -2.366 | 0.218 | 0.100 |
| Range Size | -0.129 | -0.433 | 0.166 | 0.410 |
| HFI | 0.569 | 0.190 | 0.908 | 0.004 |
| Conservation - Extinct | -1.182 | -7.880 | 5.343 | 0.794 |
| Conservation - LC | -0.542 | -2.054 | 0.879 | 0.450 |
| Conservation - Threatened | -0.931 | -2.843 | 0.886 | 0.300 |
| Biome - Grasslands | -1.255 | -2.317 | -0.091 | 0.026 |
| Biome - Mangroves | -1.510 | -7.890 | 4.615 | 0.692 |
| Biome - Med. | -0.577 | -2.678 | 1.193 | 0.556 |
| Biome - Marine | -0.788 | -2.768 | 1.194 | 0.456 |
| Biome - Taiga/Tundra | -0.411 | -1.724 | 1.069 | 0.558 |
| Biome - Temperates | -1.301 | -2.524 | -0.177 | 0.018 |
| Biome - Tropics | -1.000 | -2.123 | 0.034 | 0.070 |
| Nest structure - scrape/none | 1.644 | 0.135 | 2.931 | 0.022 |
| Nest structure - cup | -0.211 | -1.559 | 1.183 | 0.764 |
| Nest structure - platform | 1.339 | -0.171 | 2.796 | 0.072 |
| Nest structure - dome | 0.363 | -1.194 | 1.665 | 0.604 |
| Nest structure - excavation | -0.878 | -3.184 | 1.617 | 0.464 |
| Nest location - artificial | 1.143 | 0.368 | 1.916 | 0.004 |
| Nest location - ground/water | -0.311 | -1.236 | 0.629 | 0.482 |
| Nest location - cavity | -0.402 | -1.366 | 0.813 | 0.476 |
| Nest location - elevated rock | 0.214 | -0.736 | 1.142 | 0.648 |
| Nest location - vegetation | 0.221 | -0.749 | 1.280 | 0.718 |

**Figure 3:**
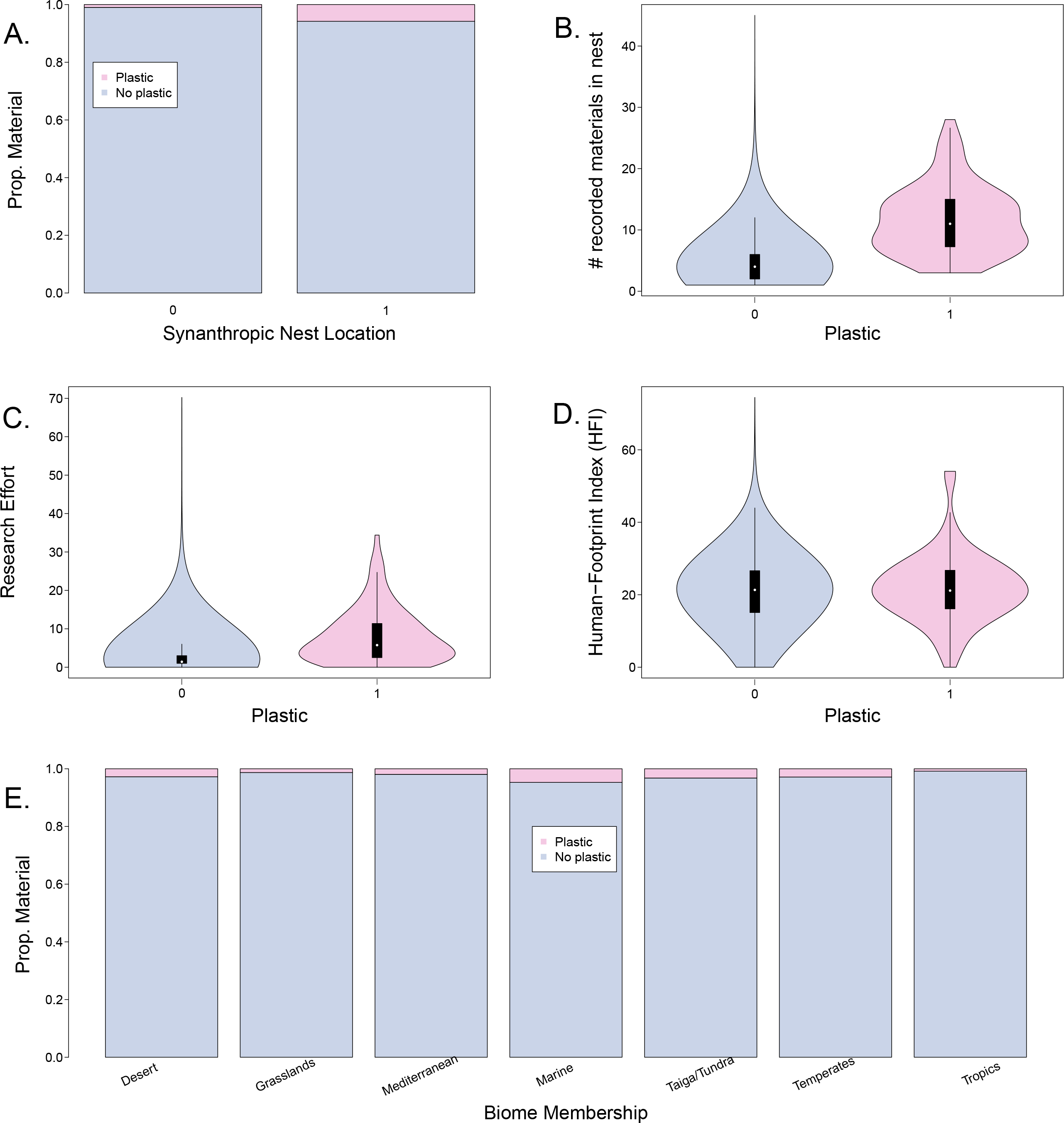
Ecological and synanthropic correlates of plastic nest material use. Correlations, uncorrected for phylogenetic signal or covariance with any other predictor, are presented at the species level between species that are (pink) and are not (purple) known to use anthropogenic nest material and A. species that are and are not known to nest in synanthropic locations, B. the number of nest materials recorded for each species, C. ‘research effort’, here estimated by the number of papers indexed by the Web of Science about that species, D. the average human-footprint index for that species, and E. assigned biome membership. For the purposes of displaying panel E, species that breed predominantly in mangroves have been omitted. All correlations shown are statistically meaningful within a phylogenetically-corrected, multivariate framework (see Table 2). Prop. = proportion.

We found no evidence that relative brain size, a potential proxy of cognitive performance and neophilia, correlates with either anthropogenic or plastic material use (anthropogenic: *β* = 1.081, *pMCMC* = 0.126, Table S3; plastic: *β* = 0.117, *pMCMC* = 0.848, Table S4).

## Discussion

We have demonstrated that bird species that incorporate plastic and anthropogenic materials into their nests are more likely to be found in human-modified landscapes, are more likely to nest in synanthropic (human-modified) locations, and incorporate more types of material into their nests, compared with species that are not known to use plastic/anthropogenic nest materials. We also demonstrate variation in anthropogenic and plastic material use between major breeding biomes; for example, anthropogenic nest material use was highest in deserts and the lowest in the tropics.

Within our models, species that are found in heavily human-modified landscapes (have a higher human-footprint index, HFI) and/or that nest in human-modified locations (e.g. nest-boxes, telephone poles, roofs, etc.) are more likely to incorporate both anthropogenic material in general and plastic material specifically into their nests. This indicates that the inclusion of anthropogenic and plastic material into nests is potentially related to the availability of these materials (Breen et al., 2021; Hansell, 2000). Our interspecific comparative data also confirm previous intraspecific correlations between anthropogenic nest material use and either HFI (e.g., Jagiello et al., 2019) or other measures of human proximity (e.g., Antczak et al., 2010; Townsend & Barker, 2014; Wang et al., 2009). This correlation between anthropogenic nest material use and tolerance to human habitation modification does not, however, seem to extend to a general high tolerance of a wide variety of ecological niches, as there is no relationship between anthropogenic/plastic material use and range size. In particular, this result suggests that species that use anthropogenic/plastic material are not necessarily generalists, which might have important conservation implications.

The suggested relationship between anthropogenic material use and material availability is further bolstered by the associations we found between anthropogenic/plastic material use and breeding biome. Anthropogenic nest material use is most prevalent in desert regions, where other types of nest material might be scarce, and is rare in tropical forests, which typically contain high amounts of biomass and somewhat lower amounts of human modification. This interpretation would accord with the data from individual bird populations, which show that nest material use is constrained by material availability (Alvarez et al., 2013; Briggs & Deeming, 2016), and that searching for nest material is energetically costly (Mainwaring & Hartley, 2013; Surgey et al., 2012). Potentially, this result could suggest that the incorporation of anthropogenic material may allow birds to expand their ranges into areas were nest materials are scarce, such as deserts, thus potentially suggests a beneficial effect of the ability to incorporate anthropogenic materials (see, e.g., Pon & Pereyra, 2021, for a study of anthropogenic nest material use in kelp gulls). The fact that there is no overall correlation between anthropogenic material use and range size, however, would indicate that such a advantage would be limited, perhaps only to certain ecological contexts.

Our analyses also suggest a potential secondary driver of anthropogenic nest material use: behavioural flexibility. Across our large sample of species, species recorded as using a higher number of nest materials – which may be due to, e.g., fewer constraints on said materials for properties such as thermal insulation (Windsor et al., 2013) or sexual signalling (Sergio et al., 2011), or due to higher levels of neophilia (Greenberg & Mettke-Hofmann, 2001) – are more likely to use both anthropogenic material generally and plastic material specifically. A species that builds nests using a wider range of materials might thus be more likely to effectively incorporate anthropogenic materials, especially those with properties not easily replicated in nature. This relationship is apparently independent of our proxy for research effort, though caution is warranted, as the number of English-language research articles indexed by Web of Science under one of potentially several synonymous scientific names does not of course fully encapsulate the total human knowledge about a species’ nest material use. Moreover, the total number of Web of Science articles about a species is an imperfect correlate of the total English-language knowledge of a species’ breeding biology; comparative studies of smaller taxonomic groups might consider more targeted proxies of reproductive research effort. Intriguingly, however, the relationship between anthropogenic nest material use and the number of recorded materials is also independent of inter-specific variation in relative brain size, a trait which correlates with neophilia and success in novel (although not necessarily urban) environments (Lefebvre et al., 2004; Sol et al., 2014; Sol et al., 2002). While this finding might in part be a consequence of the smaller sample size of the brain size models, the lack of correlation suggests, unsurprisingly, an imperfect relationship between recorded material use, neophilia, and brain size across the world’s birds.

Other than the tendency for species that use materials solely for lining (but do not excavate cavities or construct walled nests, the ‘scrape/none’ category of nest structure) to be more likely to incorporate plastic into these linings, we observed no differences in anthropogenic/plastic material use among species that build different nest types, nor among species with different body masses. Gathering materials is considered to be energetically costly (Collias & Collias, 2014; Hansell, 2000; Mainwaring & Hartley, 2013), with different nest designs potentially bearing different costs, and the effects of these costs expected to vary allometrically with body size. If anthropogenic items were a non-preferred material, they might be expected to appear more frequently in more material-heavy nests (i.e., in domes or cups instead of scrapes) or in the nests of species less able to pay metabolic costs (i.e., of smaller birds); we find no such pattern. This may in part reflect the diversity of physical properties of these ‘anthropogenic’ materials, as the costs and benefits of building a nest containing, e.g., nails (as documented in the familiar chat, *Oenanthe familiaris*) might be substantially different from those of building a nest containing, e.g., string.; an analysis examining the relevant material properties of these anthropogenic materials might be able to detect a clearer relationship between material use and energetic costs.

Perhaps controversially, in light of the increasing body of literature indicating harmful effects of the inclusion of anthropogenic materials into bird nests (Antczak et al., 2010; Carbó-Ramírez et al., 2015; O’Hanlon et al., 2021; Suárez-Rodríguez & Macías Garcia, 2014; Suárez-Rodríguez et al., 2017), we found no correlation between the incorporation of anthropogenic/plastic materials and conservation status. This may indicate that anthropogenic material use is not necessarily a current major threat to threatened and endangered species. Given the limitations in our understanding of the potential benefits and harms of anthropogenic nest material use, and given the strong correlation between research effort and the probability of detecting anthropogenic nest material use in our data, however, we urge caution in over-interpreting this result.

As human modifications to the natural world proliferate, we will also see animals increasingly attempt to respond behaviourally to these changes. Whether a species is able to react in the short-term to these habitat modifications, and whether these responses are ultimately adaptive, is an important question in understanding and mitigating the effects of the Anthropocene (Mainwaring et al., 2017). Our demonstration that one specific response, i.e., the inclusion of anthropogenic materials into bird nests, is apparently a by-product of both material availability and nest material flexibility thus underscores the importance of understanding the ecological and evolutionary origins of traits related to these behavioural consequences of human habitat modifications.

## Data Availability

The data supporting this work has been included for the reviewers’ convenience as supplementary material. Data and code will be publicly uploaded upon manuscript acceptance.

## Conflict of Interest

The authors declare no potential sources of conflicts of interest.

## Notes

### Competing Interest Statement

The authors have declared no competing interest.

